# Small Molecule Modulation of Microbiome Ecosystem: A Systems Pharmacology Perspective

**DOI:** 10.1101/2020.03.23.003285

**Authors:** Qiao Liu, Bohyun Lee, Lei Xie

## Abstract

An increasing body of evidence suggests that microbes are not only strongly associated with many human diseases but also responsible for the efficacy, resistance, and toxicity of drugs. Small-molecule drugs which can precisely fine-tune the microbial ecosystem on the basis of individual patients may revolutionize biomedicine. However, emerging endeavors in small-molecule microbiome drug discovery continue to follow a conventional “one-drug-one-target-one-disease” process. It is often insufficient and less successful in tackling complex systematic diseases. A systematic pharmacology approach that intervenes multiple interacting pathogenic species in the microbiome, could offer an attractive alternative solution. Advances in the Human Microbiome Project have provided numerous genomics data to study microbial interactions in the complex microbiome community. Integrating microbiome data with chemical genomics and other biological information enables us to delineate the landscape for the small molecule modulation of the human microbiome network. In this paper, we construct a disease-centric signed microbe-microbe interaction network using metabolite information of microbes and curated microbe effects on human health from published work. We develop a Signed Random Walk with Restart algorithm for the accurate prediction of pathogenic and commensal species. With a survey on the druggable and evolutionary space of microbe proteins, we find that 8-10% of them can be targeted by existing drugs or drug-like chemicals and that 25% of them have homologs to human proteins. We also demonstrate that drugs for diabetes are enriched in the potential inhibitors that target pathogenic microbe without affecting the commensal microbe, thus can be repurposed to modulate the microbiome ecosystem. We further show that periplasmic and cellular outer membrane proteins are overrepresented in the potential drug targets set in pathogenic microbe, but not in the commensal microbe. The systematic studies of polypharmacological landscape of the microbiome network may open a new avenue for the small-molecule drug discovery of microbiome.

**Author Summary:** As one of the most abundant components in human bodies, the microbiome has an extensive impact on human health. Pathogenic-microbes have become emerging potential therapeutic targets. Small-molecule drugs that only intervene in the growth of a specific pathogenic microbe without considering the interacting dynamics of the microbiome community may disrupt the ecosystem homeostasis, thus can cause drug side effect or prompt drug resistance. To discover novel drugs for safe and effective microbe-targeting therapeutics, a systematic approach is needed to fine-tune the microbiome ecosystem. To this end, we built a disease-centric signed microbe-microbe interaction network which accurately predicts the pathogenic or commensal effect of microbe on human health. Based on annotated and predicted pathogens and commensal species, we performed a systematic survey on therapeutic space and target landscape of existing drugs for modulating the microbiome ecosystem. Enrichment analysis on potential microbe-targeting drugs shows that drugs for diabetes could be repurposed to maintain the healthy state of microbiome. Furthermore, periplasmic and cellular outer membrane proteins are overrepresented in the potential drug targets of pathogenic-microbes, but not in proteins that perturb commensal-microbes. Our study may open a new avenue for the small molecule drug discovery of microbiome.

## Introduction

As the most abundant organism, symbiotic microbiome biomasses in human body sites are as rich as the human somatic cells [1]. Traditional culture-based or non-culture-based methods only detect limited groups of microbes, restricting our scopes on a comprehensive understanding of the entire microbial community. Advances in high throughput sequencing technology substantially enhance our powers to characterize the microbial community. Up to date, thousands of microbe genomes have been sequenced [2]. These large scale sequencing data collected have driven forward a myriad of intriguing researches, including finding microbiome biomarkers [3, 4], investigating their association with diseases [5, 6], and uncovering the dynamicity of microbial community [7, 8].

Many human genome biomarkers have been identified and utilized in disease diagnosis and precise treatment [9]. For example, small molecular drug discovery on individual cancer biomarkers has led to successful therapies [9]. Microbiome, as the “forgotten organ” of humans, contains more numerous and more diverse genomic information than the human genome. It has been shown that small molecule drugs, like antibiotics, relieve bacterial infection symptoms by controlling the overgrowth of pathogens [10]. However, many microbe species have developed antibiotics resistance mechanism, especially to several widely used drugs [11, 12]. This raises the requirement for new drugs targeting on microbes. The most direct way of drug discovery via microbe targeting is to identify drug candidates that modulate or disrupt targeted microbe proteins. The concern that comes with drug intervention treatment is their adverse effect [13, 14]. Drug intervention causes microbiota compositional change. The current view believes that microbiota homeostasis is a crucial healthy feature of our “forgotten organ”[15]. Elimination or diminution of healthy commensal microbes draws dysbiosis in our body site ecologically, and then causes symptoms like diarrhea and nausea [13]. Thus drugs minimizing side effects on other symbiotic microbes are desired.

Current studies further reveal that the microbial community is associated with a large variety of human phenotypes and diseases [5, 6]. As the microbes co-evolved with humans, many of them form mutualistic interaction with the host. Microbiota on colon mucus degrades macromolecules to metabolites, which can be digested by human colonocytes [16]. The microbes can also assist human immune system development [17]. Commensal species in human skins and colon sites expressed anti-microbial peptides (AMP) to protect against pathogenic species, like Staphylococcus aureus [18]. *Akkermansia emuciniphila* improves cancer immunotherapy efficiency [19]. Intervention on microbial community can potentially cause outgrowth of pathogenic species or suppression of commensal species, disrupting the homeostasis between microbiome and host cells and leading to diseases, including obesity [20], allergy [21], type 1 diabetes (T1D) [22] and type 2 diabetes(T2D) [23], inflammatory bowel disease (IBD) [24], rheumatoid arthritis (RA) [25], autism [26] and cancer [27]. For instance, T1D studies have shown that the abundance of Bacteroides in patient group is higher than that in the control group [22].

Symbiotic microbiota in the human body is an ecosystem. Hundreds of microbes coexist in different human body sites. The transition from healthy state to disease state often results from the disruption of microbiome system. Thus fine-tuning the microbiome ecosystem to maintain or to restore its healthy state could be effective strategy for the prevention and treatment of diseases. Existing efforts in microbiome therapeutics are dominated by probiotics or prebiotics. Despite their success in treating sudden dysbiosis like *C. difficile* infection, phenotypic responses of probiotics and prebiotics are less predictable because they change the entire microbial taxa in an unspecific way. For chronic degenerative diseases, it is needed to precisely fine-tune microbial metabolomics on the basis of individual patients. To this end, small-molecule drugs offer new opportunities and has emerged as a new frontier for microbiome drug discovery and precision medicine [28]. However, emerging endeavors in small-molecule microbiome drug discovery continue to follow a conventional “one-drug-one-target-one-disease” process. It is often insufficient and less successful in tackling complex systematic diseases. Systems pharmacology, which aims to modulate multiple targets of interacting microbiome-microbiome network, could be a potentially powerful approach to microbiome drug discovery. In order to realize systems pharmacology of microbiome, many unanswered questions remain: if there are distinct communities in the interspecies interaction network between the pathogenic microbiomes and commensal microbiomes? if we can target multiple pathogenic microbiomes, at the same time not inhibit commensal microbiomes? what the chemical space is in which the chemical compounds will inhibit pathogenic microbiome interactions but not disturb commensal microbiome? We attempt to address these questions in this perspective.

A microbe-microbe interaction network is needed for the microbiome systems pharmacology. There are many approaches to construct a microbe-microbe network based on existing evidence. The most common method is to infer this network based on microbiome occurrence abundance in the host [29, 30]. Due to the appearance of advanced sequencing technology and metagenomics, getting microbiome abundance directly from human body sites becomes available. Studies inferring the microbe-microbe interaction network is based on the abundance correlation between microbes, such as Spearman or Pearson correlation methods [31-33]. However, the microbe occurrence abundances are mostly compositional. It is still a big challenge to get absolute abundance. Plenty of research is dedicated to overcoming this problem [34, 35]. Another network inference approach is to detect microbe interactions using longitudinal data. The network generated using this method is directed and can demonstrate more various interaction types [36, 37]. However, it still suffers many issues, such as hard to get biological meaning and the requirement of optimization of sampling strategies [38, 39].

Here we constructed a disease-centric gut microbial community network by inferring microbe-microbe relationship from their metabolites input and output profiles [40], annotating microbe node in the network on their effects on human health by manually literature review, and predicting potential pathogenic and commensal microbiomes using a new signed Random Walk with Restart algorithm. We performed a survey on current knowledge of drugs-targets interactions and found a significant number of genes that could potentially be drug targets in each microbe. Our analyses suggested that large amount of genes have homologs to existing drug targets. Potential drugs, which target on pathogenic microbes with fewer side effects on the commensal microbes, can be identified. We also identified a list of potential protein targets in the pathogenic microbe and not in the commensal microbe. Our analyses took into account of the inherent factors of how drugs affect microbe growth and how microbiome interplays with each other at the molecular level. This application is not limited to the exemplar analysis performed here. It also shows the potential to filter out small molecular drugs targeting distinct pathogenic microbes based on specific patient microbiome signature. The systematic studies of polypharmacological landscape of microbiome network may open a new avenue for the small molecule drug discovery of microbiome.

## Results

### A novel disease-centric microbe-microbe interaction network

We here proposed a new microbe-microbe interaction network, which is inferred from each microbe’s metabolite consumption and production profile. Microbes affect each other through different mechanisms. 1) They have negative effects on each other through competing for the same metabolite resources (Figure 1A). 2) One microbe can have positive effects on others through cross-feeding (Figure 1A). 3) They can affect other microbes positively or negatively by alternating their living environment, like the change of pH. 4) They could also form predator-prey relationships. We can well characterize the first two relationships between microbes through inferring an interaction network using microbes’ metabolite consumption and production profiles (Figure 1B). To be specific, the extent of negative relationship is calculated as the Jaccard similarity of two microbes metabolite consumption profiles (Figure 2A). Intuitively, the more metabolites two microbes consume in common, the higher the negative effect they have on each other. On the other side, the positive effect is due to the cross-feeding relationship. The extent of positive effect is calculated as the Jaccard similarity of one microbe’s production profile and the other’s consumption profile (Figure 2B). It is worth mentioning that a positive relationship between the two microbes is not symmetrical. Finally, the extents of positive effect and negative effect are summarized to generate the final edge weights.

**Figure 1.**
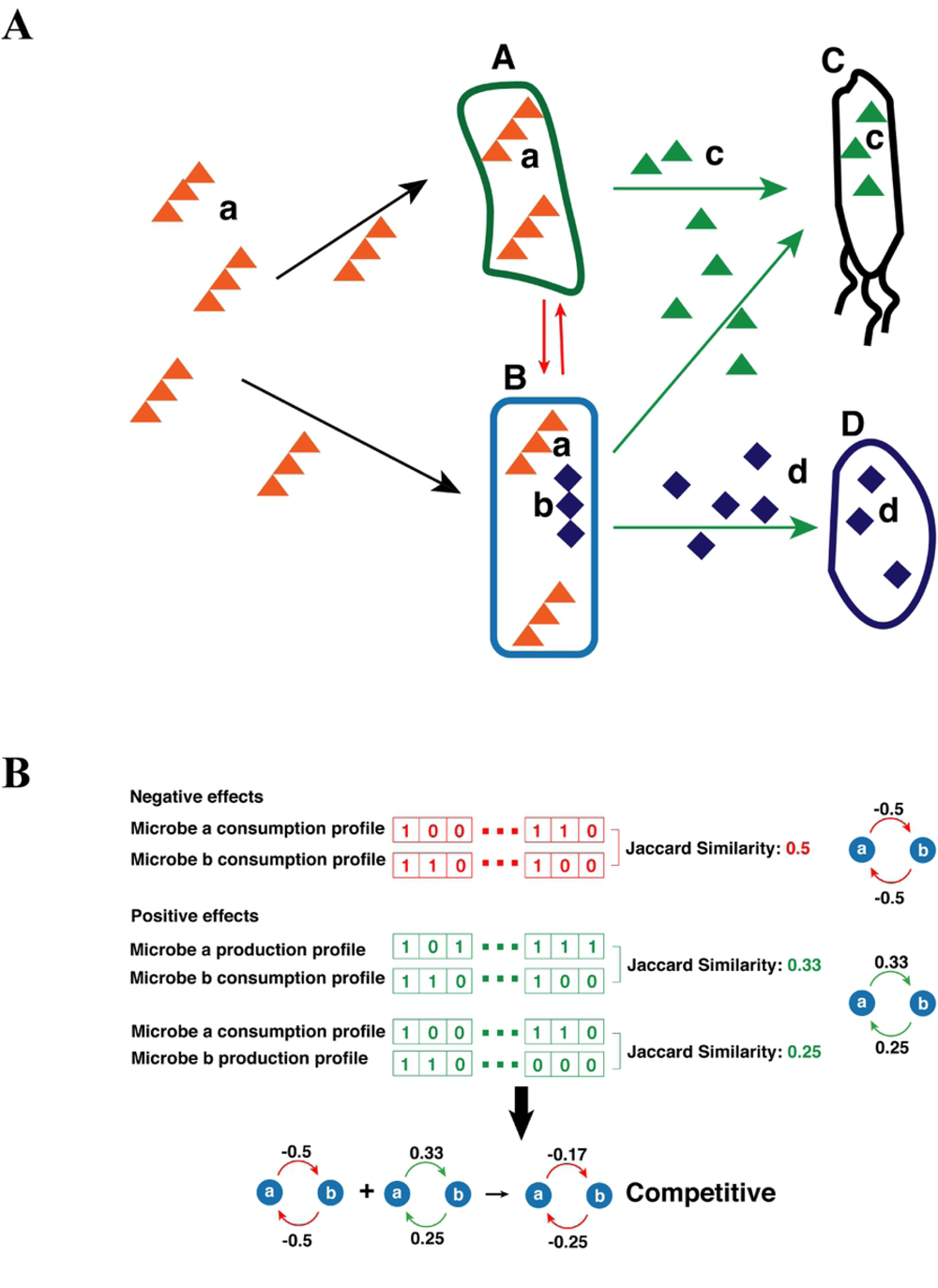
Illustration of relationships between microbiomes. A) Microbe **A** and **B** compete for metabolite **a** and has a negative effect on each other. This negative relationship is shown with a red arrow. Microbe **A** and microbe **C** have a cross-feeding relationship. Microbe **A** can degrade macromolecule a into metabolite **c**, which can be taken by microbe **C**. Cross-feedings also exist between microbe **B** and **C**, and between microbe **B** and **D** through metabolite **c** and **d**, respectively. B) An example for the calculation of the relationship between two microbes. Negative effects are calculated as the Jaccard similarity between microbes consumption profiles. The positive effect is calculated as the Jaccard similarity of one microbe’s consumption profile and another’s production profile. The final effect is the aggregation of the negative effect and the positive effect.

**Figure 2.**
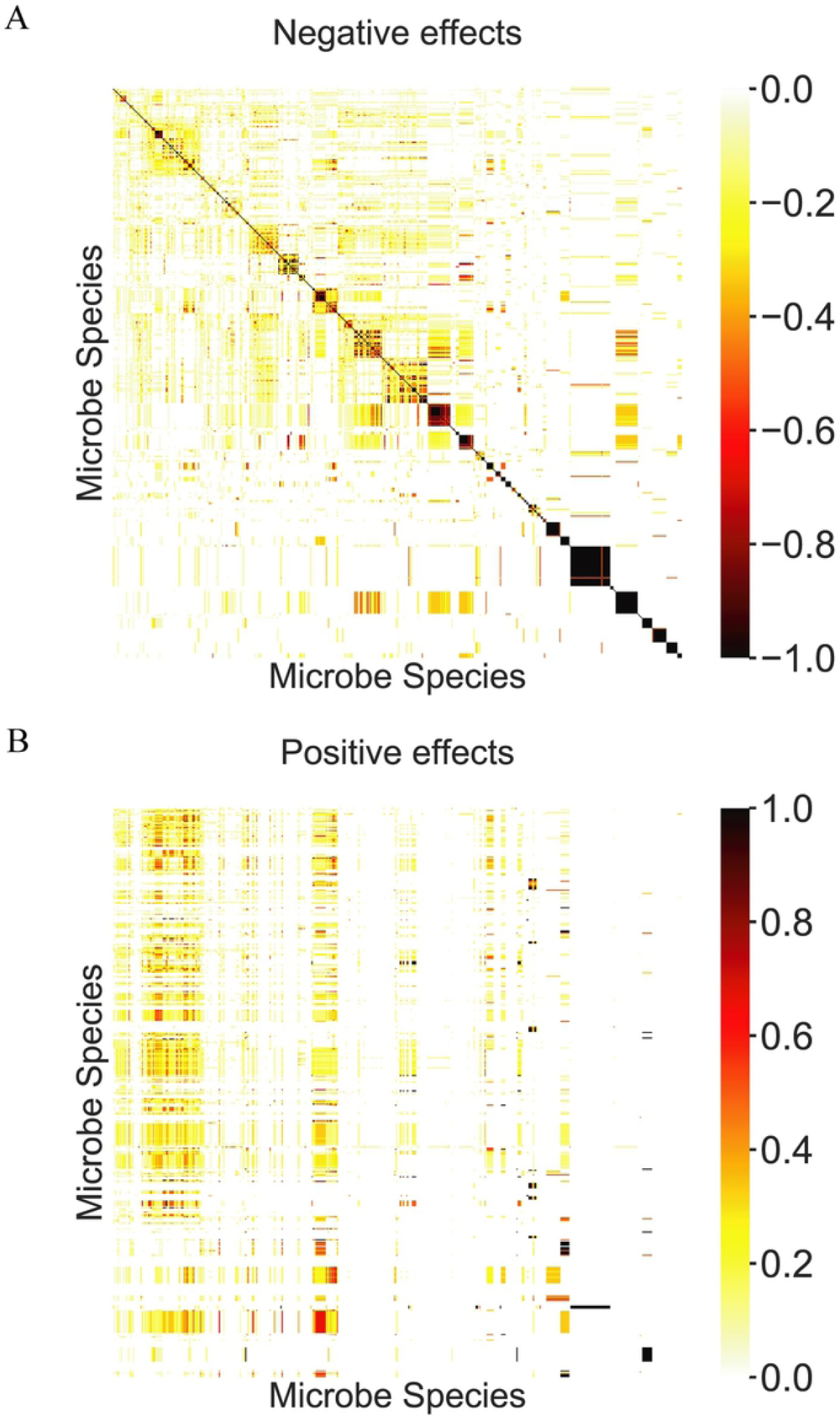
The jaccard similarity among microbes’ metabolite consumption and production profiles. A) microbe-microbe negative effect score matrix computed as the Jaccard similarity of their metabolite consumption profiles. B) microbe-microbe positive effect score matrix computed as the Jaccard similarity of one microbiome’s metabolite consumption profile and another’s metabolite production profile.

The microbe’s metabolite consumption and production profiles were curated elsewhere [40]. We include 512 microbe nodes into our network (Figure 3A). This network has the following merits: 1) The graph is directed. Therefore it can thoroughly represent various types of microbe-microbe relationships. The relationships are not limited to competition (++) and mutualism(--), where microbes can positively affect each other and negatively affect each other in both directions (Figure 3C). “+” or “-” indicates that the microbe has a positive or negative effect on the other in one direction, respectively. It can also represent more diverse relationships, including the commensalism (+ 0), parasitism (+ -) and amenalism (−0). 0 here indicates that no relationship is found in a specific direction. 2) It is biologically meaningful and straightforward to interpret. 3) This microbe-microbe interaction network can avoid the problem in the construction of microbiome network based on the abundance correlation of microbiomes, such that the correlation is sensitive to the data compositionality and is affected by low-abundance [41, 42]. 4) It can be integrated with additional networks that are derived from other information (e.g. environmental factors) into a more sophisticated heterogeneous network analysis framework.

**Figure 3.**
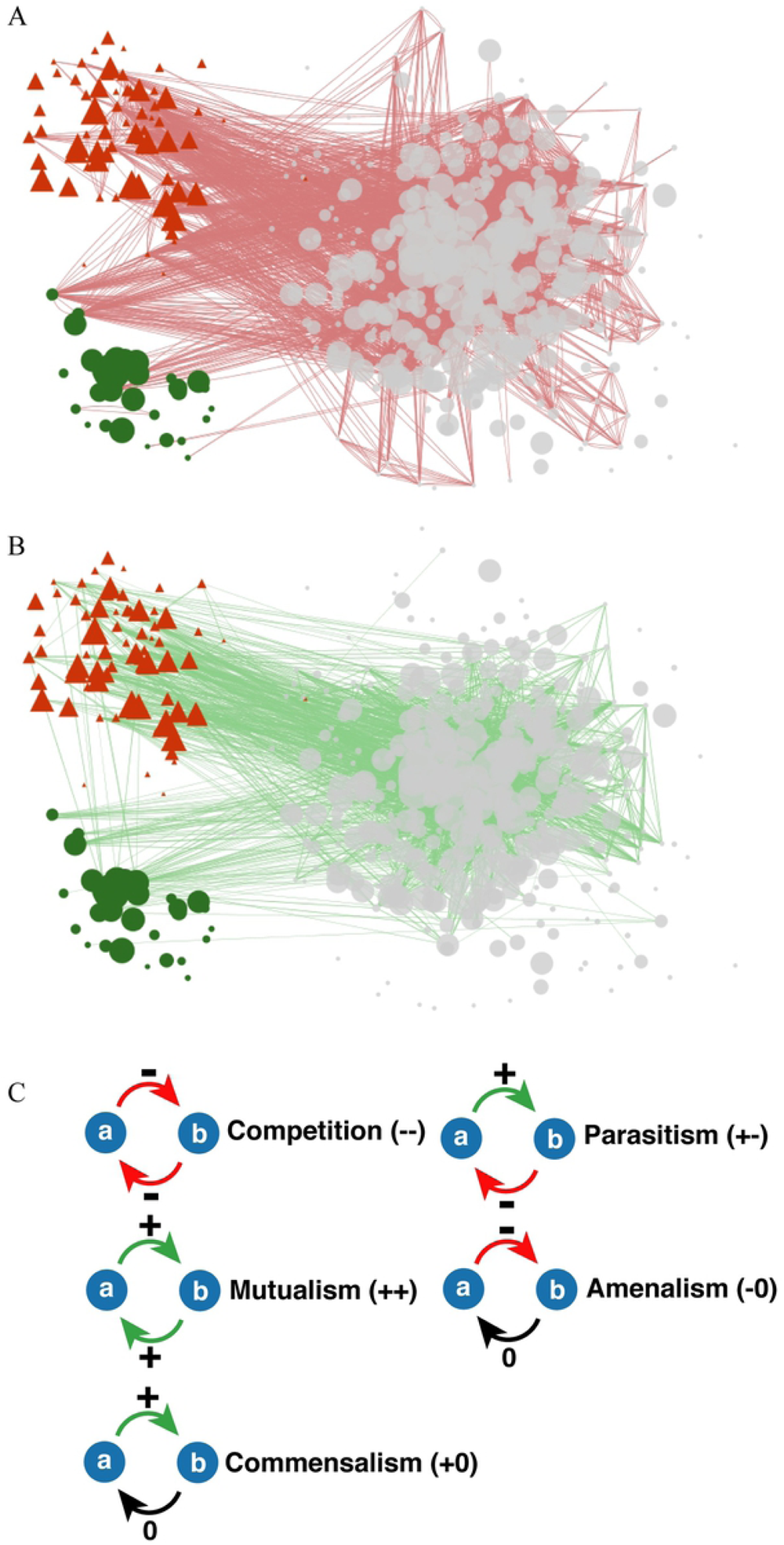
Illustration of Microbe-microbe network. A) Negative relationships (red edges) between microbes. Only the edges with weight lower than −0.7 are shown for simplicity. B) Positive relationships (green edges) between microbes. Only the edges with a weight higher than 0.5 are shown for simplicity. C) 5 relationships between microbes, competition (−-), mutualism (++), commensalism(+0), parasitism (+-) and amenalism(−0) are shown. –, + and 0 denotes a negative effect, a positive effect, and no effect, respectively. pathogenic and commensal microbes curated through literature review are labeled in red and green, respectively. Other microbes are labeled in gray.

As the “forgotten organ” of the human body, the microbiome has various effects on human health. We manually curated the microbes’ influence on human health through literature review. Out of 513 microbes in our network, we curated 72 microbes having pathogenic effects on human health and 32 microbes having commensal effects on human health (Supplementary Table S1 and Supplementary Table S2). The rest microbe influences on human health are not determined. The reasons are either lack of evidence or hard to determine the influences. Some species has different strains which have different influence on human health [43]. The microbe effects on human health are also integrated into an interaction network. The commensal microbe nodes have node weighted as “+1”, while the pathogenic microbe nodes have node weighted as “-1”. Other nodes are left unlabeled. The following analysis was performed using this network.

### Microbe effect annotation with Signed Random Walk with Restart

Because most microbe effects on human health are unknown, we developed a graph mining strategy to infer their effects based on annotated network. Using a Signed Random Walk with Restart (SRWR) model, each unannotated microbial species was treated as a node with an unknown weight, and tested for how it was influenced by the neighboring nodes in the network through corporation (positive signed edge) and competition (negative signed edge). The advantage of incorporating SRWR model into our analysis was on the fact that the network recognizes both cooperative relationships as well as the competitive relationships, which resembles the true nature of the microbial ecology. The premise of our analysis is that “friend” of “friend” or “enemy” of “enemy” will be “friend”, and “friend” of “enemy” or “enemy” of “friend” will be “enemy”.

Our SRWR simulation yielded the prediction of 143 positive nodes (potentially commensal) and 265 negative nodes (potentially pathogenic) for 418 total species without annotations associated with human health (Supplementary material Table S3). To assess the accuracy of our predictions, we use the curated data set of the 32 commensal and 72 potential pathogenic bacterial species that affects human health as a benchmark. We obtained confusion matrix with an average F1 score of 0.905. Specifically, the prediction of positive nodes yielded the precision of 0.780 and the recall of 1.000, while the prediction of negative nodes yielded the precision of 1.000 and the recall of 0.875.

### Survey on microbe proteins druggability and structural predictability

We define a microbe protein to be druggable if drugs or chemicals can target itself or its homolog. To have a comprehensive view of the microbiome, we included all protein sequences from 2232 microbe species collected by the Human Microbiome Project (HMP) in our study [2, 44]. Drugbank and ChEMBL databases are two of the most popular and updated drug-target interaction databases [45, 46]. Up to date, DrugBank and ChEMBL possessed more than 5000 and 15500 protein target sequences and drug information interacting with these targets. For each microbe species, we compared its protein sequences with the target sequences in each drug-target interaction database using PSI-Blast[47-49]. We found that a significant amount of proteins in each microbe species have homologs targeted by drugs or drug-like chemicals. The e-value resulting from a specific sequence search indicates the number of hits we can get by chance when we search a protein sequence against a database. From the plot of the percentage of protein with homologs in each microbe versus –log (e-value), we determined that the elbow point of curve is at which e-value is around 10e-60 (Figure 4A). With this e-value, we determined that 10% and 8% of microbe protein sequences were found to have close homologs in DrugBank and ChEMBL targets database. Besides, the structure information of protein is critical for the structure-based drug design and polypharmacology [50]. We searches for the homologs, which show high sequence similarities with microbe proteins, in Protein Data Bank archive (PDB) [51]. With e-value at 10e-60, we show that 25% of microbe proteins have close homologs in PDB (Figure 4B).

**Figure 4.**
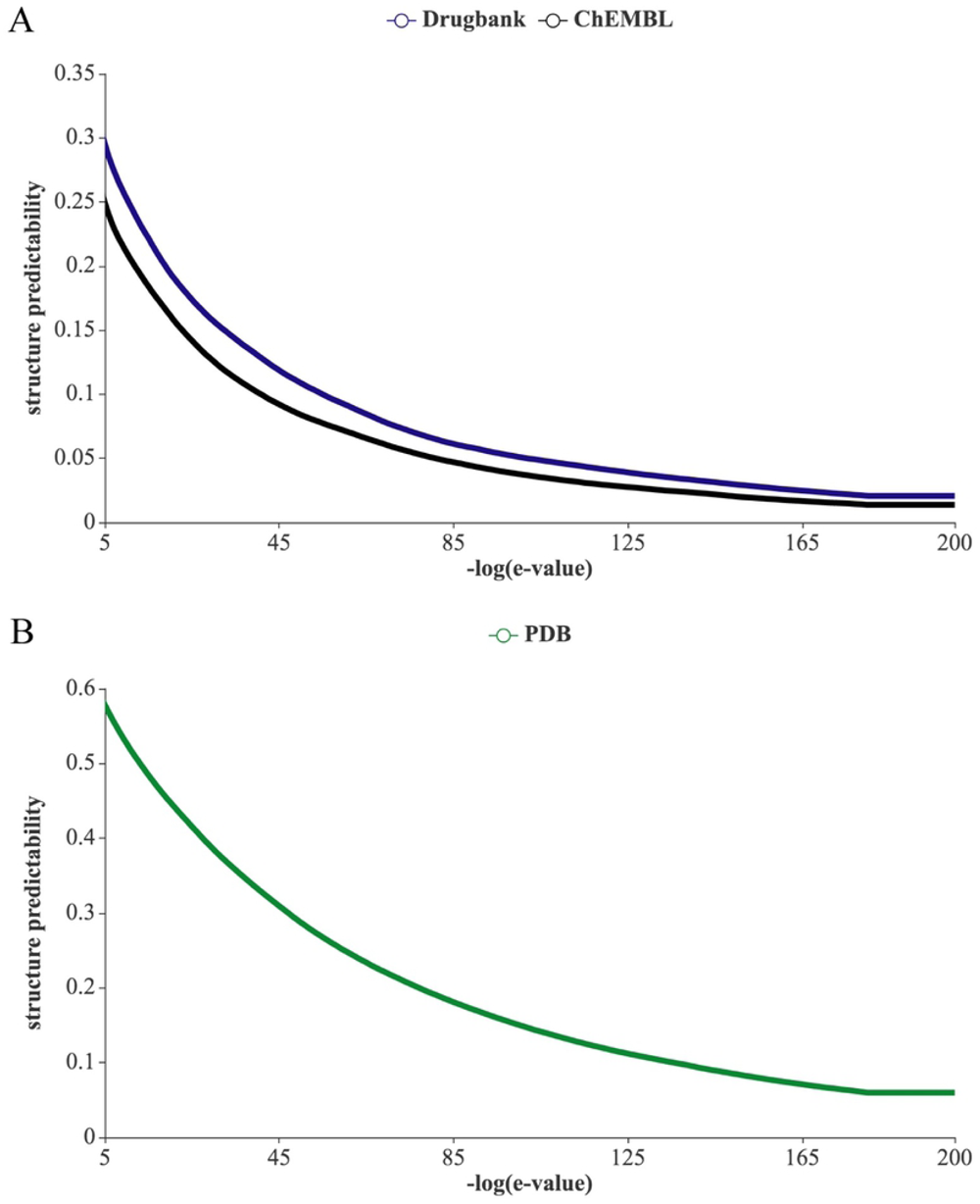
Percentage of protein targets in microbiomes that have sequence homologs in different databases, A) ChEMBL and Drugbank, and B) PDB. E-value is the criterion used in the sequence similarity search by BLAST. The lower e-value is, the closer homolog is.

### Drugs for Diabetes’ potential on drug repurposing

A successful treatment for human diseases caused by microbe infection is antibiotic intervention, which is used to control the overgrowth of one or a group of pathogenic microbes. Due to the overuse of them, many microbes have shown antibiotic resistance [11]. Another problem with some existing drugs is side effects on other symbiotic microbe species, which causes microbiome dysbiosis. Thus, disrupting pathogen interaction network by targeting multiple pathogenic microbiomes but not disturbing commensal microbiomes will be a potential powerful strategy for microbiome drug discovery. Because drug repurposing exhibits more advantages than developing a novel drug [52], we perform a computational screen on FDA approved or investigational drugs for innovative potential drugs for targeting microbes. To avoid undesirable side effects, the drugs should not affect commensal microbes proteins. With this intuition, we search for drugs that can potentially affect simultaneously multiple pathogenic microbes and avoid undesirable effect on commensal microbes.

We performed the screening on two databases: Drugbank and STITCH. Most chemicals in the Drugbank database are drugs that are FDA approved or under investigation, and most of the drug-target interactions have experimental evidence. We collected the drugs that could target proteins that are homologs of pathogenic microbes’ proteins and then excluded those targeting on homologs of commensal microbes’ proteins. Ultimately, we found 589 drugs that satisfy this constraint (Supplementary Table S4). On the other side, parts of compounds in STITCH are predicted drugs that lack experimental support. STITCH database also possesses predicted drug-targets interaction for each microbe species. Thus the screen includes both drugs and some non-drug compounds. Drug-target interactions in the STITCH database have various types, like inhibition, activation, and catalysis. We conducted more specific screening by considering each interaction type, as described in Methods. On average, one third of compounds in the STITCH database are found to have pharmaceutical usage. Finally, we found 170 drugs that appear in both STITCH screen and DrugBank screen results (Supplementary Table S5).

We then performed drugs overrepresentation analysis of these 170 drugs. The background drug list used in this analysis includes all drugs targeting microbe proteins homologs. Two analyses are conducted with two different drug classification systems, including the anatomical therapeutic chemical classification system (ATC) and the Drugbank classification system. Surprisingly, both analyses demonstrate that the drugs used in Diabetes are the statistically significantly overrepresented drugs categories (Table 1 and Table 2). Besides, it is also worth noting that the nitric oxide synthases antagonists & inhibitors are also enriched.

**Table 1.** Hypergeometric test on drugs based on drug category information in Drugbank. Only the categories with Bonferroni corrected p-value, B&H FDR adjusted p-value, or B&Y FDR adjusted p-value lower than 0.05 are shown.

| Categories | p-value | Bonferroni<br>corrected p-value | B&H FDR<br>adjusted p-value | B&Y FDR<br>adjusted p-value |
| --- | --- | --- | --- | --- |
| Nitric Oxide Synthase<br>antagonists & inhibitors | 0.00000 | 0.00012 | 0.00012 | 0.00082 |
| Urea | 0.00000 | 0.00029 | 0.00014 | 0.00094 |
| Sulfonylureas | 0.00004 | 0.01525 | 0.00508 | 0.03327 |
| Oral Hypoglycemics | 0.00007 | 0.02843 | 0.00711 | 0.04652 |
| Drugs Used in Diabetes | 0.00018 | 0.06979 | 0.01204 | 0.07882 |
| Sulfonylurea Compounds | 0.00019 | 0.07226 | 0.01204 | 0.07882 |
| Stereoisomerism | 0.00023 | 0.08839 | 0.01263 | 0.08264 |
| Aldehyde Reductase,<br>antagonists & inhibitors | 0.00041 | 0.16059 | 0.02007 | 0.13138 |
| Enzyme Inhibitors | 0.00047 | 0.18206 | 0.02023 | 0.13239 |
| Blood Glucose Lowering<br>Agents | 0.00061 | 0.23983 | 0.02398 | 0.15696 |
| Pyrazoles | 0.00135 | 0.52458 | 0.04769 | 0.31211 |

**Table 2.**
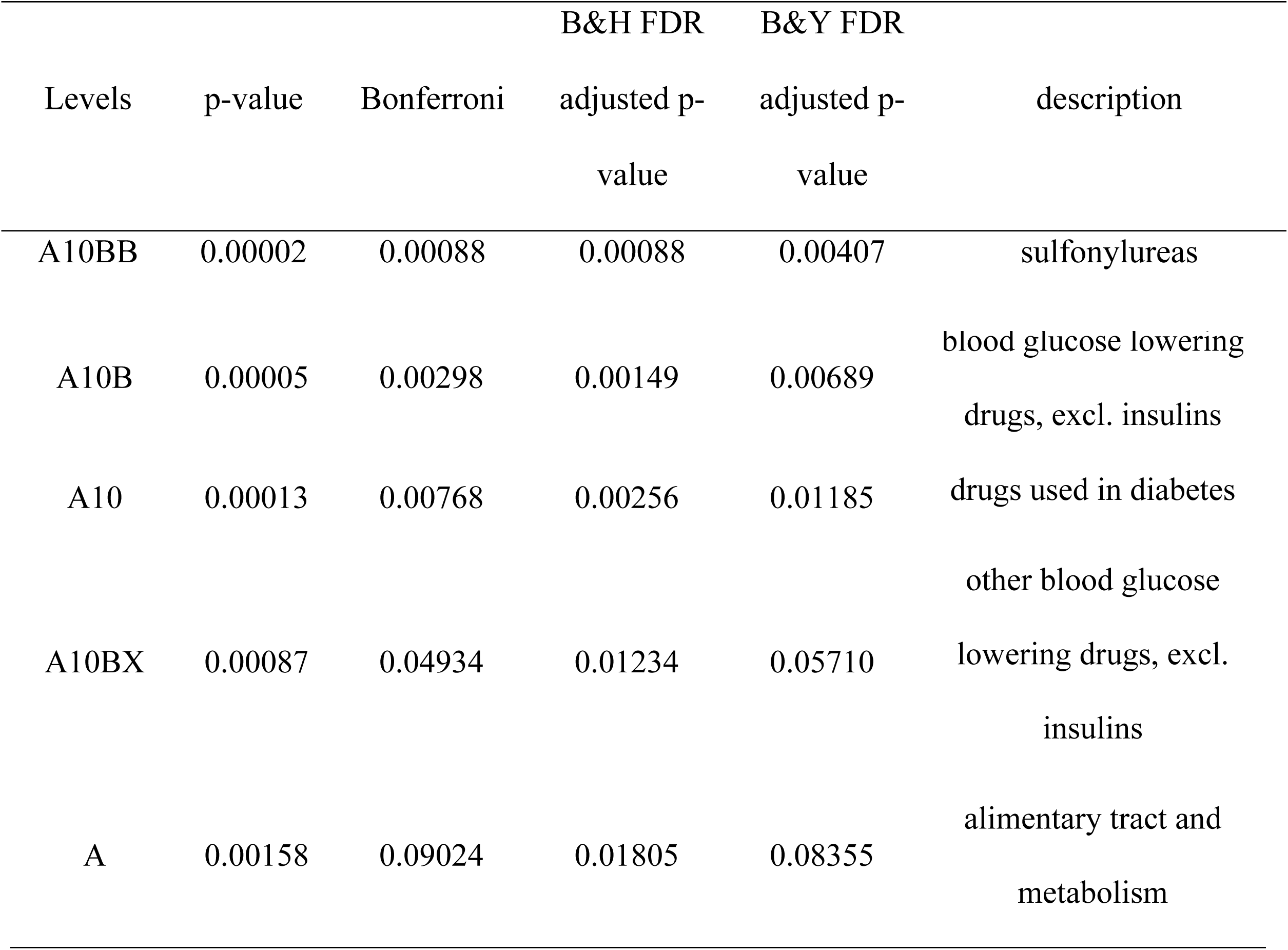
Hypergeometric test on ATC code information. Only the categories with Bonferroni corrected p-value, B&H FDR adjusted p-value, or B&Y FDR adjusted p-value lower than 0.05 are shown.

### Characterization of potential targets in pathogenic microbe proteins

We then identify targets that are homologs of pathogenic microbes’ proteins but not those of commensal microbes’ proteins. The results can assist in discerning the potential directions in drug discovery. The scope of target identification is limited to the protein targets collected in the Drugbank database. We found 462 potential proteins that are targeted by drugs (Supplementary Table S6). Functional enrichment analysis was then performed on these selected targets with DAVID [53, 54]. The background targets include all found homolog targets of microbes’ proteins that are collected by sequence search against the Drugbank database. The results show that proteins in periplasmic and cellular outer membrane are overrepresented (Table 3). The statistically significant enriched functional annotations are signal proteins and transport proteins.

**Table 3.** Results from protein targets functional annotation analysis with DAVID. Only the categories with Bonferroni corrected p-value, B&H FDR adjusted p-value, or B&H FDR adjusted p-value lower than 0.05 are shown.

| Categories | Bonferroni | B&H FDR<br>adjusted p-value | B&Y FDR<br>adjusted p-value |
| --- | --- | --- | --- |
| Periplasm | 0.00001 | 0.00001 | 0.00007 |
| Signal Peptide | 0.00010 | 0.00010 | 0.00022 |
| Topological Domain:Periplasmic | 0.00045 | 0.00022 | 0.00100 |
| GO:0009279~Cell Outer Membrane | 0.00011 | 0.00011 | 0.00104 |
| Cell Outer Membrane | 0.00028 | 0.00014 | 0.00133 |
| GO:0030288~Outer Membrane-Bounded Periplasmic Space | 0.00097 | 0.00048 | 0.00888 |
| Transmembrane Beta Strand | 0.00281 | 0.00094 | 0.01348 |
| Signal | 0.00555 | 0.00139 | 0.02669 |
| Transport | 0.01127 | 0.00226 | 0.05434 |

## Discussion

Existing small-molecule microbiome drug discovery follows conventional one-drug-one-gene-one-species paradigm [28], and focuses on preventing infection or fighting against one or a few microbes, barely considering the microbiome as an ecosystem. For examples, antibiotic has a strong influence on the composition of the microbial community and reduces the diversity of microbiome [20]. Early age onset dysbiosis, caused by antibiotic intake, potentially leads to later emergence of allergy, obesity, IBD, or other disorders. We believe systems pharmacology approaches are necessary to identify small molecule drugs for modulating the microbiome ecosystem instead of targeting a single pathogen. With the awareness of the complexity and integrity of microbiota, the reconstruction of microbiota networks is a critical step to study the microbial community and to realize systems pharmacology, and it draws increasing interests [55]. Besides occurrence abundance correlation-based methods, exploring microbes growth sources and chemical products is crucial to elucidate the mechanism of interplay between microbes.

The contemporary medical system undergoes an era of transition from traditional population based diagnosis and treatment to a more precise personalized medicine. Microbiota demonstrates high variability via developing different biogeographic signatures of human body sites [56, 57]. Small molecular drug discovery based on patient particular microbiome signatures improves and assists in generating more efficient personalized diagnosis and treatment to cure disease. Our work provides the prime landscape of small molecule drug discovery by exploring the connection between microbe’s genome and potential drugs.

Our disease-centric microbe-microbe network, constructed based on literature review and computational prediction, is still expected to improve and grow over time. Currently, the label of each node is based on species level. This method introduces ambiguity when defining each species effect on health. For instance, *E. coli*, as a prevalent commensal species in the human gut, some strains are pathogenic and even carcinogenic [43]. However, we believe that our network reflects the general effects of microbiota on health, and are useful. Firstly, most of abundant microbiomes, which we include in our network, have been well studied regarding their metabolites and effects on human health. Secondly, rare microbes effect on health inferred with the SRWR method covered the information about how they affect human health by interplaying with abundant microbiomes.

Another missing piece of information in this work is host and environmental factors. To incorporate host and environmental factors into the construction of a heterogeneous microbe-microbe interaction network will further enhance our understanding of the microbial community. Previous studies showed that environmental factors are another crucial factors determine the diversity and composition of the microbial community [58]. Gut microbiota, as the most abundant microbial community, can be affected by personal daily diet and lifestyle [59]. For example, loss of sleep could increase the ratio of Firmicutes to Bacteroidetes [60]. Microbial community is believed to harbor discrete homeostasis states and transit between different states when experiencing environmental changes, at least for skin or vaginal microbiota [61, 62]. Thus, constructing a heterogeneous human-environment-microbiome network will be an important direction for the future work.

## Methods

### Microbiome interaction network

Microbe species metabolite consumption and production information were manually curated elsewhere [40]. 513 microbe species are included in this dataset. Distribution of the number of metabolites each microbe consumes or produces, and distribution of the number of microbes each metabolite associates with are investigated (Supplementary Figure 1). We hypothesize that the final relationship between the two microbes is composed of a negative relationship (competition) and a positive relationship (corporation). The negative extent, negative_ab_, is calculated as the Jaccard similarity of metabolite consumption profile between microbe a and microbe b. 233 metabolites are investigated and are consumed by at least one microbe species. The positive extent, positive_ab_, is calculated as the Jaccard similarity of microbiome a’s metabolites consumption profile to microbiome b’s production profile. At least one microbe species produces 95 metabolites. The final microbiome interaction network is a directed graph. The edge weight between two microbes is calculated as positive_ab_ - negative_ab_.

### Signed Random Walk with Restart (SRWR)

Microbiome interaction network with selected 512 microbial species were simulated using Signed Random Walk with Restart algorithm [63]. To predict the label (pathogenic or commensal) of a unannotated microbiome species, it was initialized as a start node. For each run, initial score of 1.0 was assigned to the start node with an unknown sign, and then this score was distributed out to the neighboring nodes via edges in the network as the walk goes with random probability. Positive edge would increase the positive ranking score of the neighboring node with the balance attenuation probability of β=0.5, and the negative edge would increase the negative ranking score of the neighboring node with the probability of γ = 0.5. When the walk was complete, positive scores from known commensal species, and negative scores from known pathogenic species were summed up to predict the label for the unknown start node (supplementary table S1, S2 and S3).

### Microbe proteins druggability survey

Protein sequences of 2232 microbes were downloaded from Human Microbiome Projects (HMP) [2, 44]. Druggable target sequences were downloaded from DrugBank (www.drugbank.ca) and ChEMBL websites (www.ebi.ac.uk/chembl) [45, 46]. They were saved as fasta format and reformatted to be a Blast database using PSI-Blast tools [47-49]. Each microbe protein sequences were searched against each target sequence database to find their homologs. Biopython package was used to perform sequence search using PSI-Blast [47-49]. All sequence search results with e-value lower than 10e-4 were saved to generate a plot of percentage of protein with homologs in each microbe versus –log (e-value). Scripts used for analysis are available in https://github.com/qiaoliuhub/drug_target_analysis_on_microbiome.

### Potential drugs screening

#### Drugbank

By using the homolog targets from the sequence search, we collected the drugs that potentially target microbe proteins for each microbe. The drug-target interaction database was downloaded from DrugBank (www.drugbank.ca). All drugs that target pathogenic microbes are gathered into a candidate list, then parts of the drugs in the list are excluded if they can potentially target commensal microbes. 589 drugs in the candidate list were left after screening.

#### STITCH

STITCH database was downloaded from *http://stitch.embl.de/* [64]. STITCH database has grouped drug-target interactions based on microbe species. These drug-target interactions are also classified into different types, such as inhibition, activation, or catalysis. We focus on two interaction types: inhibition and activation. Our primary purpose is to screen for FDA approved or investigational drugs, so we excluded non-drugs compounds. We utilized the STITCH drug ID information, which is also the same with PubChem compound ID, to retrieve the pharmaceutical function information from the PubChem database by using PUG REST API and E-utilities tools [65]. We performed the following screen: 1) compounds that activate targets in pathogenic microbes but not activate targets in commensal microbes (134 drugs). 2) compounds that activate targets in pathogenic microbes but not inhibit targets in commensal microbes (431 drugs). 3) compounds that inhibit targets in pathogenic microbes but not inhibit targets in commensal microbes (185 drugs). 4) compounds that inhibit targets in pathogenic microbes but not activate targets in commensal microbes (1325 drugs) (Supplementary Table S7).

#### The intersection of Drugbank screening and STITCH screening result

The InChIKey information of all drugs found in Drugbank screening was retrieved from Drugbank full database XML file. The InChIKey information of all drugs found in STITCH screening was collected from the Pubchem website using PUG REST API and E-utilities tools. The intersection of these two drugs InChIKey list was found for later analysis.

### Overrepresentation analysis

#### Drug overrepresentation analysis

All drugs ATC code and Drugbank classification information were accumulated from Drugbank full database XML file. These two classification systems have hierarchical structures, and all categories in all levels are included. A hypergeometric test was performed on the Drugbank screened 589 drugs list. The Bonferroni correction, Benjamini & Hochberg’sHochberg’s FDR adjustment, and Benjamini & Yekutieli’sYekutieli’s FDR adjustment methods were used to adjust the p-values of these multiple comparisons. ∼3700 drugs, which are found to target at least one microbe protein homolog, are used as background drugs list (Supplementary Table S8).

### Protein targets functional enrichment analysis

462 potential protein targets were filtered out using the Drugbank target sequences database and saved as UniProt accession number. Target’s functional enrichment analysis was conducted with the database for annotation, visualization, and integrated discovery (DAVID) [53, 54]. The list of 462 potential targets’ UniProt accession numbers was uploaded to DAVID as a test gene set. Background gene set includes ∼1700 microbe protein homologs (Supplementary Table S9).

## Acknowledgements

This work was supported by Grant Number R01GM122845 from the National Institute of General Medical Sciences (NIGMS) and Grand Number R01AD057555 of National Institute of Aging of the National Institute of Health (NIH) as well as CUNY High Performance Computing Center. The funders had no role in study design, data collection and analysis, decision to publish, or preparation of the manuscript.

## Author contributions

Q.L., B.L. and L.X. designed the computational framework. Q.L. and B.L. carried out the implementation and analyzed the data. Q.L. and B.L. wrote the manuscript with input from all authors. L.X. revised the manuscript, conceived and planned the study.

## Supporting information captions

Supplementary Table S1. Microbes with pathogenic effects on human health by manually literature review.

Supplementary Table S2. Microbes with commensal effects on human health by manually literature review.

Supplementary Table S3. The Microbes with unknown effects in literature reviews and their SRWR inferred microbe effects.

Supplementary Table S4. Drug screen results using Drugbank database.

Supplementary Table S5. Drugs that are found in both drug screen results using Drugbank database and that using STITCH database.

Supplementary Table S6. Potential homolog proteins that have homologs with pathogenic microbe proteins but do not have homologs with commensal microbe proteins.

Supplementary Table S7. Drug screen results in STITCH database.

Supplementary Table S8. Background drugs list used in drug overrepresentation analysis.

Supplementary Table S9. Background targets list used in target functional annotation analysis.

Supplementary Figure 1. A) Distribution of number of metabolites each microbiome consume or produce. B) Distribution of number of microbiomes each metabolite.

## Reference

1. Sender R, Fuchs S, Milo R. Are We Really Vastly Outnumbered? Revisiting the Ratio of Bacterial to Host Cells in Humans. Cell. 2016;164(3):337–40. Epub 2016/01/30. doi:10.1016/j.cell.2016.01.013. PubMed PMID: 26824647.

2. Human Microbiome Project C. Structure, function and diversity of the healthy human microbiome. Nature. 2012;486(7402):207–14. Epub 2012/06/16. doi:10.1038/nature11234. PubMed PMID: 22699609; PubMed Central PMCID: PMCPMC3564958.

3. Rajpoot M, Sharma AK, Sharma A, Gupta GK. Understanding the microbiome: Emerging biomarkers for exploiting the microbiota for personalized medicine against cancer. Semin Cancer Biol. 2018;52(Pt 1):1–8. Epub 2018/02/10. doi:10.1016/j.semcancer.2018.02.003. PubMed PMID: 29425888.

4. Dietert RR, Silbergeld EK. Biomarkers for the 21st century: listening to the microbiome. Toxicol Sci. 2015;144(2):208–16. Epub 2015/03/22. doi:10.1093/toxsci/kfv013. PubMed PMID: 25795652.

5. Jackson MA, Verdi S, Maxan ME, Shin CM, Zierer J, Bowyer RCE, et al. Gut microbiota associations with common diseases and prescription medications in a population-based cohort. Nat Commun. 2018;9(1):2655. Epub 2018/07/10. doi:10.1038/s41467-018-05184-7. PubMed PMID: 29985401; PubMed Central PMCID: PMCPMC6037668.

6. Kho ZY, Lal SK. The Human Gut Microbiome - A Potential Controller of Wellness and Disease. Front Microbiol. 2018;9:1835. Epub 2018/08/30. doi:10.3389/fmicb.2018.01835. PubMed PMID: 30154767; PubMed Central PMCID: PMCPMC6102370.

7. Thaiss CA. Microbiome dynamics in obesity. Science. 2018;362(6417):903–4. Epub 2018/11/24. doi:10.1126/science.aav6870. PubMed PMID: 30467161.

8. Nunn KL, Forney LJ. Unraveling the Dynamics of the Human Vaginal Microbiome. Yale J Biol Med. 2016;89(3):331–7. Epub 2016/10/05. PubMed PMID: 27698617; PubMed Central PMCID: PMCPMC5045142.

9. Hoelder S, Clarke PA, Workman P. Discovery of small molecule cancer drugs: successes, challenges and opportunities. Mol Oncol. 2012;6(2):155–76. Epub 2012/03/24. doi:10.1016/j.molonc.2012.02.004. PubMed PMID: 22440008; PubMed Central PMCID: PMCPMC3476506.

10. Wilson ID, Nicholson JK. Gut microbiome interactions with drug metabolism, efficacy, and toxicity. Transl Res. 2017;179:204–22. Epub 2016/09/04. doi:10.1016/j.trsl.2016.08.002. PubMed PMID: 27591027; PubMed Central PMCID: PMCPMC5718288.

11. Zaman SB, Hussain MA, Nye R, Mehta V, Mamun KT, Hossain N. A Review on Antibiotic Resistance: Alarm Bells are Ringing. Cureus. 2017;9(6):e1403. Epub 2017/08/31. doi:10.7759/cureus.1403. PubMed PMID: 28852600; PubMed Central PMCID: PMCPMC5573035.

12. Zinner SH. Antibiotic use: present and future. New Microbiol. 2007;30(3):321–5. Epub 2007/09/07. PubMed PMID: 17802919.

13. Jernberg C, Lofmark S, Edlund C, Jansson JK. Long-term impacts of antibiotic exposure on the human intestinal microbiota. Microbiology. 2010;156(Pt 11):3216–23. Epub 2010/08/14. doi:10.1099/mic.0.040618-0. PubMed PMID: 20705661.

14. Dudek-Wicher RK, Junka A, Bartoszewicz M. The influence of antibiotics and dietary components on gut microbiota. Prz Gastroenterol. 2018;13(2):85–92. Epub 2018/07/14. doi:10.5114/pg.2018.76005. PubMed PMID: 30002765; PubMed Central PMCID: PMCPMC6040098.

15. Lloyd-Price J, Abu-Ali G, Huttenhower C. The healthy human microbiome. Genome Med. 2016;8(1):51. Epub 2016/04/29. doi:10.1186/s13073-016-0307-y. PubMed PMID: 27122046; PubMed Central PMCID: PMCPMC4848870.

16. Samuel BS, Hansen EE, Manchester JK, Coutinho PM, Henrissat B, Fulton R, et al. Genomic and metabolic adaptations of Methanobrevibacter smithii to the human gut. Proc Natl Acad Sci U S A. 2007;104(25):10643–8. Epub 2007/06/15. doi:10.1073/pnas.0704189104. PubMed PMID: 17563350; PubMed Central PMCID: PMCPMC1890564.

17. Belkaid Y, Hand TW. Role of the microbiota in immunity and inflammation. Cell. 2014;157(1):121–41. Epub 2014/04/01. doi:10.1016/j.cell.2014.03.011. PubMed PMID: 24679531; PubMed Central PMCID: PMCPMC4056765.

18. Nakatsuji T, Chen TH, Narala S, Chun KA, Two AM, Yun T, et al. Antimicrobials from human skin commensal bacteria protect against Staphylococcus aureus and are deficient in atopic dermatitis. Sci Transl Med. 2017;9(378). Epub 2017/02/24. doi:10.1126/scitranslmed.aah4680. PubMed PMID: 28228596; PubMed Central PMCID: PMCPMC5600545.

19. Routy B, Le Chatelier E, Derosa L, Duong CPM, Alou MT, Daillere R, et al. Gut microbiome influences efficacy of PD-1-based immunotherapy against epithelial tumors. Science. 2018;359(6371):91–7. Epub 2017/11/04. doi:10.1126/science.aan3706. PubMed PMID: 29097494.

20. Trasande L, Blustein J, Liu M, Corwin E, Cox LM, Blaser MJ. Infant antibiotic exposures and early-life body mass. Int J Obes (Lond). 2013;37(1):16–23. Epub 2012/08/22. doi:10.1038/ijo.2012.132. PubMed PMID: 22907693; PubMed Central PMCID: PMCPMC3798029.

21. Fujimura KE, Sitarik AR, Havstad S, Lin DL, Levan S, Fadrosh D, et al. Neonatal gut microbiota associates with childhood multisensitized atopy and T cell differentiation. Nat Med. 2016;22(10):1187–91. Epub 2016/09/13. doi:10.1038/nm.4176. PubMed PMID: 27618652; PubMed Central PMCID: PMCPMC5053876.

22. de Goffau MC, Luopajarvi K, Knip M, Ilonen J, Ruohtula T, Harkonen T, et al. Fecal microbiota composition differs between children with beta-cell autoimmunity and those without. Diabetes. 2013;62(4):1238–44. Epub 2013/01/01. doi:10.2337/db12-0526. PubMed PMID: 23274889; PubMed Central PMCID: PMCPMC3609581.

23. Qin J, Li Y, Cai Z, Li S, Zhu J, Zhang F, et al. A metagenome-wide association study of gut microbiota in type 2 diabetes. Nature. 2012;490(7418):55–60. Epub 2012/10/02. doi:10.1038/nature11450. PubMed PMID: 23023125.

24. Huttenhower C, Kostic AD, Xavier RJ. Inflammatory bowel disease as a model for translating the microbiome. Immunity. 2014;40(6):843–54. Epub 2014/06/21. doi:10.1016/j.immuni.2014.05.013. PubMed PMID: 24950204; PubMed Central PMCID: PMCPMC4135443.

25. Scher JU, Sczesnak A, Longman RS, Segata N, Ubeda C, Bielski C, et al. Expansion of intestinal Prevotella copri correlates with enhanced susceptibility to arthritis. Elife. 2013;2:e01202. Epub 2013/11/07. doi:10.7554/eLife.01202. PubMed PMID: 24192039; PubMed Central PMCID: PMCPMC3816614.

26. Kang DW, Adams JB, Gregory AC, Borody T, Chittick L, Fasano A, et al. Microbiota Transfer Therapy alters gut ecosystem and improves gastrointestinal and autism symptoms: an open-label study. Microbiome. 2017;5(1):10. Epub 2017/01/27. doi:10.1186/s40168-016-0225-PubMed PMID: 28122648; PubMed Central PMCID: PMCPMC5264285.

27. Kostic AD, Chun E, Robertson L, Glickman JN, Gallini CA, Michaud M, et al. Fusobacterium nucleatum potentiates intestinal tumorigenesis and modulates the tumor-immune microenvironment. Cell Host Microbe. 2013;14(2):207–15. Epub 2013/08/21. doi:10.1016/j.chom.2013.07.007. PubMed PMID: 23954159; PubMed Central PMCID: PMCPMC3772512.

28. Cully M. Microbiome therapeutics go small molecule. Nat Rev Drug Discov. 2019;18(8):569–72. Epub 2019/08/02. doi:10.1038/d41573-019-00122-8. PubMed PMID: 31367062.

29. Ban Y, An L, Jiang H. Investigating microbial co-occurrence patterns based on metagenomic compositional data. Bioinformatics. 2015;31(20):3322–9. Epub 2015/06/17. doi:10.1093/bioinformatics/btv364. PubMed PMID: 26079350; PubMed Central PMCID: PMCPMC4795632.

30. Berry D, Widder S. Deciphering microbial interactions and detecting keystone species with co-occurrence networks. Front Microbiol. 2014;5:219. Epub 2014/06/07. doi:10.3389/fmicb.2014.00219. PubMed PMID: 24904535; PubMed Central PMCID: PMCPMC4033041.

31. Arumugam M, Raes J, Pelletier E, Le Paslier D, Yamada T, Mende DR, et al. Enterotypes of the human gut microbiome. Nature. 2011;473(7346):174–80. Epub 2011/04/22. doi:10.1038/nature09944. PubMed PMID: 21508958; PubMed Central PMCID: PMCPMC3728647.

32. Barberan A, Bates ST, Casamayor EO, Fierer N. Using network analysis to explore co-occurrence patterns in soil microbial communities. ISME J. 2012;6(2):343–51. Epub 2011/09/09. doi:10.1038/ismej.2011.119. PubMed PMID: 21900968; PubMed Central PMCID: PMCPMC3260507.

33. Copeland JK, Yuan L, Layeghifard M, Wang PW, Guttman DS. Seasonal community succession of the phyllosphere microbiome. Mol Plant Microbe Interact. 2015;28(3):274–85. Epub 2015/02/14. doi:10.1094/MPMI-10-14-0331-FI. PubMed PMID: 25679538.

34. Li H. Microbiome, Metagenomics, and High-Dimensional Compositional Data Analysis. Annual Review of Statistics and Its Application. 2015;2(1):73–94. doi:10.1146/annurev-statistics-010814-020351.

35. Tsilimigras MC, Fodor AA. Compositional data analysis of the microbiome: fundamentals, tools, and challenges. Ann Epidemiol. 2016;26(5):330–5. Epub 2016/06/04. doi:10.1016/j.annepidem.2016.03.002. PubMed PMID: 27255738.

36. Baksi KD, Kuntal BK, Mande SS. ‘TIME’: A Web Application for Obtaining Insights into Microbial Ecology Using Longitudinal Microbiome Data. Front Microbiol. 2018;9:36. Epub 2018/02/09. doi:10.3389/fmicb.2018.00036. PubMed PMID: 29416530; PubMed Central PMCID: PMCPMC5787560.

37. Caporaso JG, Lauber CL, Costello EK, Berg-Lyons D, Gonzalez A, Stombaugh J, et al. Moving pictures of the human microbiome. Genome Biol. 2011;12(5):R50. Epub 2011/06/01. doi:10.1186/gb-2011-12-5-r50. PubMed PMID: 21624126; PubMed Central PMCID: PMCPMC3271711.

38. Gerber GK. The dynamic microbiome. FEBS Lett. 2014;588(22):4131–9. Epub 2014/03/04. doi:10.1016/j.febslet.2014.02.037. PubMed PMID: 24583074.

39. Datta MS, Sliwerska E, Gore J, Polz MF, Cordero OX. Microbial interactions lead to rapid micro-scale successions on model marine particles. Nat Commun. 2016;7:11965. Epub 2016/06/18. doi:10.1038/ncomms11965. PubMed PMID: 27311813; PubMed Central PMCID: PMCPMC4915023.

40. Sung J, Kim S, Cabatbat JJT, Jang S, Jin YS, Jung GY, et al. Global metabolic interaction network of the human gut microbiota for context-specific community-scale analysis. Nat Commun. 2017;8:15393. Epub 2017/06/07. doi:10.1038/ncomms15393. PubMed PMID: 28585563; PubMed Central PMCID: PMCPMC5467172.

41. Layeghifard M, Hwang DM, Guttman DS. Disentangling Interactions in the Microbiome: A Network Perspective. Trends Microbiol. 2017;25(3):217–28. Epub 2016/12/06. doi:10.1016/j.tim.2016.11.008. PubMed PMID: 27916383.

42. Rottjers L, Faust K. From hairballs to hypotheses-biological insights from microbial networks. FEMS Microbiol Rev. 2018;42(6):761–80. Epub 2018/08/08. doi:10.1093/femsre/fuy030. PubMed PMID: 30085090; PubMed Central PMCID: PMCPMC6199531.

43. Martinez-Medina M, Garcia-Gil LJ. Escherichia coli in chronic inflammatory bowel diseases: An update on adherent invasive Escherichia coli pathogenicity. World J Gastrointest Pathophysiol. 2014;5(3):213–27. Epub 2014/08/19. doi:10.4291/wjgp.v5.i3.213. PubMed PMID: 25133024; PubMed Central PMCID: PMCPMC4133521.

44. Human Microbiome Project C. A framework for human microbiome research. Nature. 2012;486(7402):215–21. Epub 2012/06/16. doi:10.1038/nature11209. PubMed PMID: 22699610; PubMed Central PMCID: PMCPMC3377744.

45. Wishart DS, Feunang YD, Guo AC, Lo EJ, Marcu A, Grant JR, et al. DrugBank 5.0: a major update to the DrugBank database for 2018. Nucleic Acids Res. 2018;46(D1):D1074–D82. Epub 2017/11/11. doi:10.1093/nar/gkx1037. PubMed PMID: 29126136; PubMed Central PMCID: PMCPMC5753335.

46. Gaulton A, Hersey A, Nowotka M, Bento AP, Chambers J, Mendez D, et al. The ChEMBL database in 2017. Nucleic Acids Res. 2017;45(D1):D945–D54. Epub 2016/12/03. doi:10.1093/nar/gkw1074. PubMed PMID: 27899562; PubMed Central PMCID: PMCPMC5210557.

47. Camacho C, Coulouris G, Avagyan V, Ma N, Papadopoulos J, Bealer K, et al. BLAST+: architecture and applications. BMC Bioinformatics. 2009;10:421. Epub 2009/12/17. doi:10.1186/1471-2105-10-421. PubMed PMID: 20003500; PubMed Central PMCID: PMCPMC2803857.

48. Altschul SF, Madden TL, Schaffer AA, Zhang J, Zhang Z, Miller W, et al. Gapped BLAST and PSI-BLAST: a new generation of protein database search programs. Nucleic Acids Res. 1997;25(17):3389–402. Epub 1997/09/01. doi:10.1093/nar/25.17.3389. PubMed PMID: 9254694; PubMed Central PMCID: PMCPMC146917.

49. Altschul SF, Gish W, Miller W, Myers EW, Lipman DJ. Basic local alignment search tool. J Mol Biol. 1990;215(3):403–10. Epub 1990/10/05. doi:10.1016/S0022-2836(05)80360-2. PubMed PMID: 2231712.

50. Xie L, Xie L, Kinnings SL, Bourne PE. Novel computational approaches to polypharmacology as a means to define responses to individual drugs. Annu Rev Pharmacol Toxicol. 2012;52:361–79. Epub 2011/10/25. doi:10.1146/annurev-pharmtox-010611-134630. PubMed PMID: 22017683.

51. Berman HM, Westbrook J, Feng Z, Gilliland G, Bhat TN, Weissig H, et al. The Protein Data Bank. Nucleic Acids Res. 2000;28(1):235–42. Epub 1999/12/11. doi:10.1093/nar/28.1.235. PubMed PMID: 10592235; PubMed Central PMCID: PMCPMC102472.

52. Pushpakom S, Iorio F, Eyers PA, Escott KJ, Hopper S, Wells A, et al. Drug repurposing: progress, challenges and recommendations. Nat Rev Drug Discov. 2019;18(1):41–58. Epub 2018/10/13. doi:10.1038/nrd.2018.168. PubMed PMID: 30310233.

53. Huang da W, Sherman BT, Lempicki RA. Systematic and integrative analysis of large gene lists using DAVID bioinformatics resources. Nat Protoc. 2009;4(1):44–57. Epub 2009/01/10. doi:10.1038/nprot.2008.211. PubMed PMID: 19131956.

54. Huang da W, Sherman BT, Lempicki RA. Bioinformatics enrichment tools: paths toward the comprehensive functional analysis of large gene lists. Nucleic Acids Res. 2009;37(1):1–13. Epub 2008/11/27. doi:10.1093/nar/gkn923. PubMed PMID: 19033363; PubMed Central PMCID: PMCPMC2615629.

55. Xiao Y, Angulo MT, Friedman J, Waldor MK, Weiss ST, Liu YY. Mapping the ecological networks of microbial communities. Nat Commun. 2017;8(1):2042. Epub 2017/12/13. doi:10.1038/s41467-017-02090-2. PubMed PMID: 29229902; PubMed Central PMCID: PMCPMC5725606.

56. Hannigan GD, Duhaime MB, Koutra D, Schloss PD. Biogeography and environmental conditions shape bacteriophage-bacteria networks across the human microbiome. PLoS Comput Biol. 2018;14(4):e1006099. Epub 2018/04/19. doi:10.1371/journal.pcbi.1006099. PubMed PMID: 29668682; PubMed Central PMCID: PMCPMC5927471.

57. Oh J, Byrd AL, Deming C, Conlan S, Program NCS, Kong HH, et al. Biogeography and individuality shape function in the human skin metagenome. Nature. 2014;514(7520):59–64. Epub 2014/10/04. doi:10.1038/nature13786. PubMed PMID: 25279917; PubMed Central PMCID: PMCPMC4185404.

58. xxxiii Goodrich JK, Waters JL, Poole AC, Sutter JL, Koren O, Blekhman R, et al. Human genetics shape the gut microbiome. Cell. 2014;159(4):789–99. Epub 2014/11/25. doi:10.1016/j.cell.2014.09.053. PubMed PMID: 25417156; PubMed Central PMCID: PMCPMC4255478.

59. Gilbert JA, Blaser MJ, Caporaso JG, Jansson JK, Lynch SV, Knight R. Current understanding of the human microbiome. Nat Med. 2018;24(4):392–400. Epub 2018/04/11. doi:10.1038/nm.4517. PubMed PMID: 29634682.

60. Benedict C, Vogel H, Jonas W, Woting A, Blaut M, Schurmann A, et al. Gut microbiota and glucometabolic alterations in response to recurrent partial sleep deprivation in normal-weight young individuals. Mol Metab. 2016;5(12):1175–86. Epub 2016/12/03. doi:10.1016/j.molmet.2016.10.003. PubMed PMID: 27900260; PubMed Central PMCID: PMCPMC5123208.

61. Ravel J, Gajer P, Abdo Z, Schneider GM, Koenig SS, McCulle SL, et al. Vaginal microbiome of reproductive-age women. Proc Natl Acad Sci U S A. 2011;108 Suppl 1:4680–7. Epub 2010/06/11. doi:10.1073/pnas.1002611107. PubMed PMID: 20534435; PubMed Central PMCID: PMCPMC3063603.

62. DiGiulio DB, Callahan BJ, McMurdie PJ, Costello EK, Lyell DJ, Robaczewska A, et al. Temporal and spatial variation of the human microbiota during pregnancy. Proc Natl Acad Sci U S A. 2015;112(35):11060–5. Epub 2015/08/19. doi:10.1073/pnas.1502875112. PubMed PMID: 26283357; PubMed Central PMCID: PMCPMC4568272.

63. Jung J, Jin W, Sael L, Kang U, editors. Personalized Ranking in Signed Networks Using Signed Random Walk with Restart. 2016 IEEE 16th International Conference on Data Mining (ICDM); 2016 12-15 Dec. 2016.

64. Szklarczyk D, Santos A, von Mering C, Jensen LJ, Bork P, Kuhn M. STITCH 5: augmenting protein-chemical interaction networks with tissue and affinity data. Nucleic Acids Res. 2016;44(D1):D380–4. Epub 2015/11/22. doi:10.1093/nar/gkv1277. PubMed PMID: 26590256; PubMed Central PMCID: PMCPMC4702904.

65. Kim S, Chen J, Cheng T, Gindulyte A, He J, He S, et al. PubChem 2019 update: improved access to chemical data. Nucleic Acids Res. 2019;47(D1):D1102–D9. Epub 2018/10/30. doi:10.1093/nar/gky1033. PubMed PMID: 30371825; PubMed Central PMCID: PMCPMC6324075.

